# Adaptive Kinase Signalling Enables Escape from Small Molecule Inhibition in Glioblastoma

**DOI:** 10.1101/2025.10.28.684838

**Authors:** Mike van Heumen, Linde Hoosemans, Jolanda Piepers, Sander S.M. Rensen, Kim R. Kampen, Ann Hoeben, Marc A. Vooijs

**Affiliations:** Department of Radiation Oncology (MAASTRO), GROW School for Oncology and Reproduction, Maastricht University Medical Center+, Maastricht, Netherlands; Department of Medical Oncology, GROW School for Oncology and Reproduction, Maastricht University Medical Center+, Maastricht, Netherlands; Department of Surgery, NUTRIM Institute for Nutrition and Translational Research in Metabolism, Maastricht University Medical Center+, Maastricht, Netherlands

**Keywords:** Glioblastoma, small molecule inhibitor, therapy resistance, combination therapy, kinase activity, signalling pathway

## Abstract

Patients with glioblastoma (GB) have a median survival of 15 months. Despite an intensive treatment schedule with resection, radio- and chemotherapy, recurrence is inevitable. A significant challenge to overcome GB treatment resistance involves intratumoral heterogeneity, characterized by molecular, phenotypic, and clinical distinctive GB subtypes. Different small molecule inhibitors (SMI) have been designed to inhibit signalling proteins in oncogenic driver pathways in GB. However, SMIs have been unsuccessful in improving patient outcomes. Here, we investigate whether crosstalk between signalling pathways and signalling pathway redundancy are responsible for single-agent resistance using a primary patient-derived glioblastoma organoid (PGO) platform.

This study used nine FDA-approved small-molecule inhibitors, based on their ability to cross the blood-brain barrier, targeting key GB driver genes. Although inhibition of downstream effector proteins reduced cell viability more effectively (IC50 70nM-1μM) than inhibiting upstream membrane-bound tyrosine kinase receptors (IC50 1-15μM), remaining cell proliferation was seen in all six PGOs. To uncover resistance mechanisms, we analysed phosphokinase activity following monotherapy in various PGOs and identified compensatory pathway activation, leading to the discovery of effective small-molecule inhibitor combinations, most notably CHIR99021 (GSK-3 inhibitor) with trametinib (MEK inhibitor).

In conclusion, our findings highlight the potential of combination therapy targeting compensatory pathways to overcome single-agent resistance in GB, emphasizing the utility of patient-derived glioblastoma organoids as a platform for personalized therapeutic development.

## 1. Introduction

Glioblastoma (GB) is the most common malignant primary brain tumour in adults (1). Standard-of-care includes maximal surgical resection followed by chemoradiation and adjuvant chemotherapy using temozolomide (TMZ) (2, 3). However, survival benefits from this multimodal approach are minimal, and recurrence is inevitable. No life-prolonging treatments exist in the recurrent setting, where median overall survival is less than six months (4). Novel treatment options are required to improve clinical outcomes. Despite multiple clinical trials, including trials with immune checkpoint inhibitors, no survival benefit for GB patients could be obtained (5, 6).

An important challenge in GB treatment is intratumoral heterogeneity, which involves the presence of phenotypically distinct GB subclones. These are marked by their own driver mutations e.g. neural progenitor-like (NP-like), oligodendrocyte-progenitor-like (OPC-like), astrocyte-like (AC-like), and mesenchymal-like (MES-like) states, which impede the effectiveness of single agents (7).

The AC-like subtype is characterised by mutated or amplified epidermal growth factor receptor (*EGFR)* and OPC-like tumours are characterised by amplified platelet derived growth factor receptor (*PDGFR)*, which both activate the downstream RAS/RAF pathway (7-9). This RAS/RAF pathway is also frequently upregulated in mesenchymal-like tumours by amplified hepatocyte growth factor receptor (*c-MET*). Moreover, mutations, deletions and reduced expression of the tumour suppressor gene phosphatase and tensin homolog *(PTEN)* in classical, proneural and mesenchymal tumours lead to AKT activation through phosphatidylinositol (3,4,5)-triphosphate (*PIP*_*3*_) amplification. Lastly, in classical, proneural and mesenchymal GB, mouse double minute 2 homolog *(MDM2/4)* is frequently amplified, leading to increased ubiquitination and degradation of TP53 in these tumours (8). These distinct cellular states often coexist within a single tumor, highlighting GB’s profound intratumoral heterogeneity and the challenge of developing pan-GB effective therapies. Upon treatment, GB can undergo further genomic alterations through clonal selection, allowing resistant subclones to persist and ultimately drive recurrence and treatment resistance (7, 10-12).

Various small molecule inhibitors (SMIs) have been shown to induce reduced cell viability in GB cell models *in vitro*, such as afatinib (EGFR inhibitor), abemaciclib (CDK4/CDK6 inhibitor), buparlisib (PI3K inhibitor), vistusertib (mTOR inhibitor), cediranib (VEGFR/c-KIT/PDGFRß inhibitor) and costunolide (TERT inhibitor) (11-13). However, these SMIs have not improved overall survival in clinical trials, leading to an unchanged standard of care for patients with a newly diagnosed GB since 2005 (2, 6, 14).

Substantial evidence from *in-vitro* models and clinical studies shows both intrinsic and acquired resistance of cancers to SMIs. For instance, in non-small cell lung cancer, patients often develop resistance to EGFR inhibitors within the first year of treatment, frequently due to compensatory signalling such as c-MET amplification and mutations in the RAS/RAF and PI3K/Akt pathways (15-18). These resistance mechanisms can be countered by c-MET or MEK co-inhibition, highlighting the value of combinatorial therapy (17-19).

In GB, simultaneous activation of multiple pathways may underlie poor responses to TKIs, and combined inhibition of multiple receptor tyrosine kinases (RTKs) has been shown to more effectively reduce cell viability in GB cell models (20). In addition to having simultaneously activating pathways, GB tumours can also activate compensatory pathways to bypass the effects of a single SMI. Important bypass mechanisms studied in GB include the crosstalk between the PI3K/mTOR pathway and MEK/ERK pathway and the upregulation of MET or AXL after treatment with an EGFR inhibitor (21-25). Combinatorial therapies targeting these bypass mechanisms have been studied preclinically (26, 27). However, clinical trials have not yet led to improved survival and new approved treatments (28, 29). This is likely due to dose-limiting toxicity to healthy tissue, limited local drug efficacy and the use of an unstratified patient population (28, 30). Hence, there is a clinical need to identify rational and effective combinational therapies in biomarker driven individualized clinical trials.

As a first step, we investigated the effects of single-agent and combination therapies using approved blood-brain barrier-penetrant SMIs targeting the most prominent oncogenic drivers in GB, using patient-derived glioblastoma organoids (PGO). These GB organoids resemble the phenotypic and genotypic features of the parental tumour and maintain their intratumoral genetic and phenotypic heterogeneity in culture (31).

We demonstrate that each PGO responds differently to selected SMIs, corresponding to a unique compensatory phosphokinase profile. Rational drug combinations targeting these compensatory kinase regulations reduced cell viability in a patient-specific manner.

## 2. Material and Methods

### 2.1 Patient recruitment

Tumor tissue from patients diagnosed with IDH1/2 wild-type GB was collected and processed according to a protocol approved by the Medical Ethics Committee Zuyderland (METC-Z; study number 17-T-101). The study was registered under clinical trial identifiers NCT04865315 and NCT04868396. Eligible participants were adults (≥18 years) scheduled for surgical resection of a lesion suspected to be GBM based on magnetic resonance imaging findings. Written informed consent was obtained from all patients prior to surgery.

Tumor specimens were obtained at Maastricht University Medical Center+ (MUMC+) and Zuyderland Medical Center (Heerlen). Immediately after surgical removal, tissue samples were processed to generate PGOs following a previously published protocol (31). Additional samples (1919, 3565, and 3128) were generously provided by collaborating research groups.

### 2.1 Cell Culture

Six patient-derived glioblastoma organoid lines (PGOs) were obtained as described before (31, 32). In brief, GB samples obtained from patients were subjected to mechanical and enzymatic dissociation to generate single-cell suspensions, which were subsequently resuspended in Cultrex Basement Membrane Extract (R&D Systems; #3433-010-01) to establish PGOs. PGOs were cultured in Neurobasal Medium complete (NBMc) containing Neurobasal Medium (ThermoFisher #21103049), vitamin B-27 supplement minus vitamin A (ThermoFisher #12587010), Antibiotic-Antimycotic (ThermoFisher #15240062), L-Glutamine (ThermoFisher #25030081), AmphotericinB (ThermoFisher #15290018), 10ng/mL recombinant human EGF (ThermoFisher) and 10 ng/mL recombinant human basic FGF (ThermoFisher). PGOs were cultured at 37°C in a 95% humidified atmosphere containing 5% CO_2_. All cell lines were regularly tested for mycoplasma contamination using qPCR with primers targeting S16 and were confirmed to be negative.

### 2.2 Cell viability Assays

For the cell viability assays, PGOs were dissociated into single cells using TripLE (ThermoFisher #12605036) to enable standardized seeding. Cells were plated in quadruplicates at a density of 1000 cells/well in a 384-wells plate (Greiner; #781098) coated with 1:100 diluted Cultrex Basement Membrane Extract in Neurobasal medium to allow for re-aggregation into 3D multicellular clusters. The cells were incubated for 24h before being subjected to treatment.

Cells were treated with different concentrations of drugs, including abemaciclib, afatinib, APR-246, axitinib, buparlisib, sunitinib, and trametinib (range 80 nM – 50 µM) for 7 days using a TECAN D300e drug dispenser. CellTiterGlo® 3D Luminescent Cell Viability Assay (Promega; #G968) was performed per manufacturer’s instructions to assess cell viability. The luminescence signal was measured using the Spectramax iD3 microplate reader.

### 2.3 Combinatorial treatments

Cell viability assays were performed as described above or via the use of Alamar Blue (ThermoFisher, Cat#DAL1100) following the manufacturer’s protocol, with a 3-hour incubation period. SMIs were diluted in a 5-point dilution series, which were selected to best represent the dose-response range of each inhibitor based on single agent efficacy. Combinations were created with TECAN D300E software in which two or three different inhibitors were combined in all concentrations.

### 2.4 Long term exposure assays

For long-term drug exposure experiments, two GB cell lines, U87 and U1242, were dissociated into single cells using Trypsin (ThermoFisher, Cat#15400054). Cells were seeded in triplicate at a density of 500 cells per well in 24-wells plates (Greiner, Cat#662160) and allowed to adhere for 24 hours. Treatments were administered on days 1, 4, 8, and 11 using trametinib (1.25–25 nM) and/or CHIR99021, a GSK3β inhibitor (0.5–2 µM). All compounds were prepared in fresh Dulbecco’s Modified Eagle Medium (DMEM; Sigma-Aldrich, Cat#D6429) supplemented with 10% fetal bovine serum (FBS; Gibco, Cat#10500-064). DMSO concentration was normalized to 1%.

At day 10 or 14, cell viability was assessed using Alamar Blue (ThermoFisher, Cat#DAL1100) following the manufacturer’s protocol, with a 3-hour incubation period. Subsequently, cells were washed with PBS, fixed with 4% formaldehyde (ThermoFisher, Cat#43368.9M), and stained with 0.1% crystal violet (Sigma-Aldrich, Cat#10034810980). Plates were imaged, and staining intensity was analysed using Image Studio Lite (LI-COR Biosciences). For additional quantification, 0.5 mL of 10% acetic acid (Sigma-Aldrich, Cat#A6283) was added to each well to solubilize the dye, and absorbance was measured at 595 nm using a SpectraMax iD3 plate reader.

For patient-derived glioma organoids (PGOs 031 and 009), 50,000 cells per well were seeded in NBMc, in triplicate, in 24-wells plates. Following a 24-hour incubation, cells were treated on days 1, 4, 8, and 11 with trametinib (1.25–25 nM) and/or CHIR99021 (0.25–1 µM), dissolved in NBMc medium containing 1% DMSO. Fresh medium was only added at day 8. Cell viability was evaluated on day 15 using flow cytometry (Cytek Guava easyCyte) after staining with propidium iodide (PI).

### 2.5 Protein extraction and Western Blot analysis

Cells were cultured via standardised cell-culture protocols and seeded on 10-cm dishes with 2.5·10^6^ cells per dish at day 1. Treatment consisting of a SMI or DMSO (vehicle-control) was added on day 3 at 1 µM, and lysates were prepared on day 4. Cells were scraped, washed, and incubated with RIPA buffer plus protease and phosphatase inhibitor cocktail (Thermofisher; #78440) for 20 minutes on ice. Subsequently, cells were snap-frozen in liquid nitrogen and thawed three times before centrifuging at 13.000g for 15min. Protein concentrations were quantified using Bradford analysis (Bio-Rad), and protein samples were prepared in 1x Laemmli loading buffer. Equal amounts of proteins were loaded on Tris-HCL SDS-PAGE gels and transferred onto nitrocellulose membranes. Membranes were blocked in 0.05% Tween20 in TBS (TBS-T) with 5% dried skimmed milk (Marvel). Membranes were incubated overnight with primary antibodies (Supplementary Table 1). After washing with tris-buffered saline with tween (TBS-T), membranes were incubated for 1 hour at room temperature with either anti-rabbit (CST; #7074) or anti-mouse (CST; #7076) horseradish peroxidase (HRP) conjugated secondary antibodies. Amersham ECL Prime Western Blotting Detection Reagent (GE Healthcare) was used to visualise the proteins as described by the manufacturer in the Azure C600 immunoblot imager. A protein ladder (ThermoFisher: #26619) was run and transferred alongside the samples to determine molecular weight. Blot images were subsequently overlaid with the corresponding ladder to verify the kDa of the detected protein bands. WB to assess on-target activity of drugs were performed at least twice on independent experiments and showed consistent results.

### 2.6 Phosphokinase array assays

Lysates (100 µg) from PGOs treated with 1µM abemaciclib, buparlisib, trametinib or DMSO for 7 days were incubated with the Proteome Profiler Human Phospho-Kinase Array Kit (R&D Systems; # ARY003C) and Proteome Profiler Human Phospho-RTK Array Kit (R&D Systems; #ARY001B) according to the manufacturer’s instructions. The chemiluminescent signal was revealed using the Azure C600 immunoblot imager. The signalling intensity of the phospho-proteins was analysed with array software (Quick Spots, OptimEyes). For each membrane, the background (e.g. pixel intensity of negative controls) was subtracted from each signal intensity. Duplicate intensities exceeding a pixel intensity of 500, with a difference of less than 100, were included in further analysis.

Protein array proteomics data was transformed into relative values to their respective DMSO controls. Data (matrix input data) was uploaded into SRplot (PMID: 37943830) to generate Principal Component Analysis (PCA) and a Euclidean hierarchical cluster heatmap. The divergent cluster #2 was further analysed with Reactome pathway analysis.

### 2.8 Gene expression profiling

Gene expression data for GSK-3ß was analysed using GEPIA (Gene Expression Profiling Interactive Analysis), a web tool that integrates data from The Cancer Genome Atlas (TCGA) and Genotype-Tissue Expression (GTEx) databases. (33) We selected TCGA datasets relevant to GB, and differential gene expression was evaluated using the tool’s default parameters. Expression levels were compared between tumour and normal tissue samples, and statistical significance was assessed using a p-value threshold of 0.05. Survival analyses were performed to examine the association between gene expression levels and overall survival.

### 2.9 Statistical Analysis

Unless otherwise stated, all experiments were performed using three independent biological replicates. Data are presented as mean values, with error bars indicating the variability between biological replicates. Western blot analyses performed exclusively to confirm on-target drug activity were conducted once and are therefore presented as representative experiments. Graphs were created using GraphPad Prism (v.9.2.0) by normalising the luminescence values of the treatment conditions to the DMSO control. IC50s were calculated using “log(inhibitor) vs. response -- Variable slope (four parameters)” in GraphPad Prism (v.9.2.0).

The SynergyFinder+ tool (https://synergyfinder.org/) was used to identify synergistic combinations using the Bliss score for combination treatment. (34) A Bliss score>10 was considered synergistic, -10 to 10 was considered additive, and <-10 was considered antagonistic.

## 3 Results

### 3.1 Drug screen

Nine small-molecule inhibitors (Table S4) were chosen for their potential to target prevalent mutations in GB. These inhibitors are FDA-approved and can cross the blood-brain-barrier, except for APR-246, a drug restoring TP53’s tumour suppressive function. Each inhibitor was tested on six genetically diverse patient-derived glioma organoids (PGOs) at varying concentrations (80 nM to 50 µM). The degree of reduction in cell viability differed among the SMI (Figure 1 and S1). Five out of nine inhibitors obtained an IC50 at concentrations under 2µM across all PGOs, including abemaciclib (CDK4/6 inhibitor), afatinib (EGFR inhibitor), axitinib (VEGFR, PDGFRβ and c-Kit inhibitor), buparlisib (PI3K inhibitor), and trametinib (MEK inhibitor) (Figure 1A-C and S1). Notably, inhibitors targeting downstream effector proteins, such as abemaciclib, trametinib, and buparlisib , showed lower IC50s compared to SMIs targeting upstream membrane-bound tyrosine kinase receptors, like afatinib and axitinib . No correlation was observed between the variations in IC50 across the PGOs and their mutational status (Figure 1D and Table S2/3) (31).

**Fig 1.**
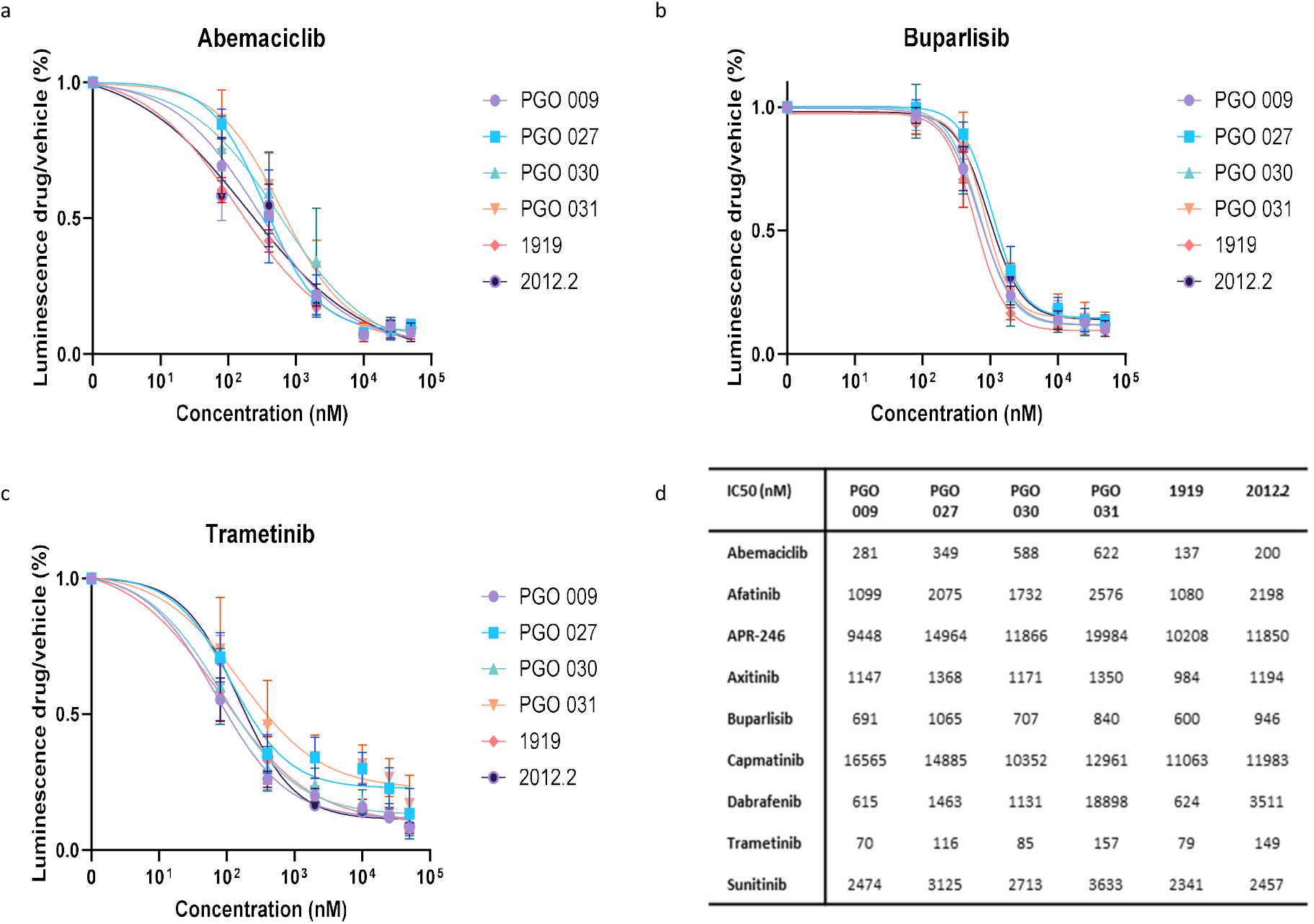
Treatment sensitivity of patient-derived glioblastoma organoids (PGOs) to selected small-molecule inhibitors (SMIs). Cell viability was assessed using CellTiter-Glo after 7 days of drug exposure and is presented as dose response curves for each PGO following treatment with the indicated inhibitors: (a) abemaciclib (CDK4/CDK6 inhibitor), (b) buparlisib (PI3K inhibitor), or (c) trametinib (MEK inhibitor). Data represent the mean ± SD of n = 3 biological replicates. (d) Inhibitory concentration 50% (IC50) values in nM for each PGO and the respective SMIs.

### 3.2 On-target efficacy

Next, we assessed the on-target inhibition of each SMI in all PGOs. Treatment with abemaciclib, afatinib, buparlisib, and trametinib (1 µM) reduced the activation of their corresponding downstream targets (Figure 2A-C and S2). The degree of inhibition varied among PGOs for certain inhibitors. Specifically, phosphorylation of AKT was most markedly suppressed in PGO 009 following buparlisib treatment, whereas inhibition of phosphorylated ERK upon afatinib exposure was more pronounced in PGOs 027, 031, and 2012.2 (Figure B and S2). Moreover, afatinib treatment resulted in a reduction in initial target activity, as shown by decreased phospho-EGFR levels (Figure S2). However, total EGFR levels were increased after treatment with afatinib . Abemaciclib reduced total Rb levels and completely blocked phospho-Rb levels in all PGOs (Figure 2A). Following trametinib treatment, phospho-MEK (Ser217/221) was hyperphosphorylated compared to the vehicle control (Figure 2C). Subsequently, no downstream phospho-ERK (Tyr202/204) was detectable in trametinib-treated samples .

**Fig 2.**
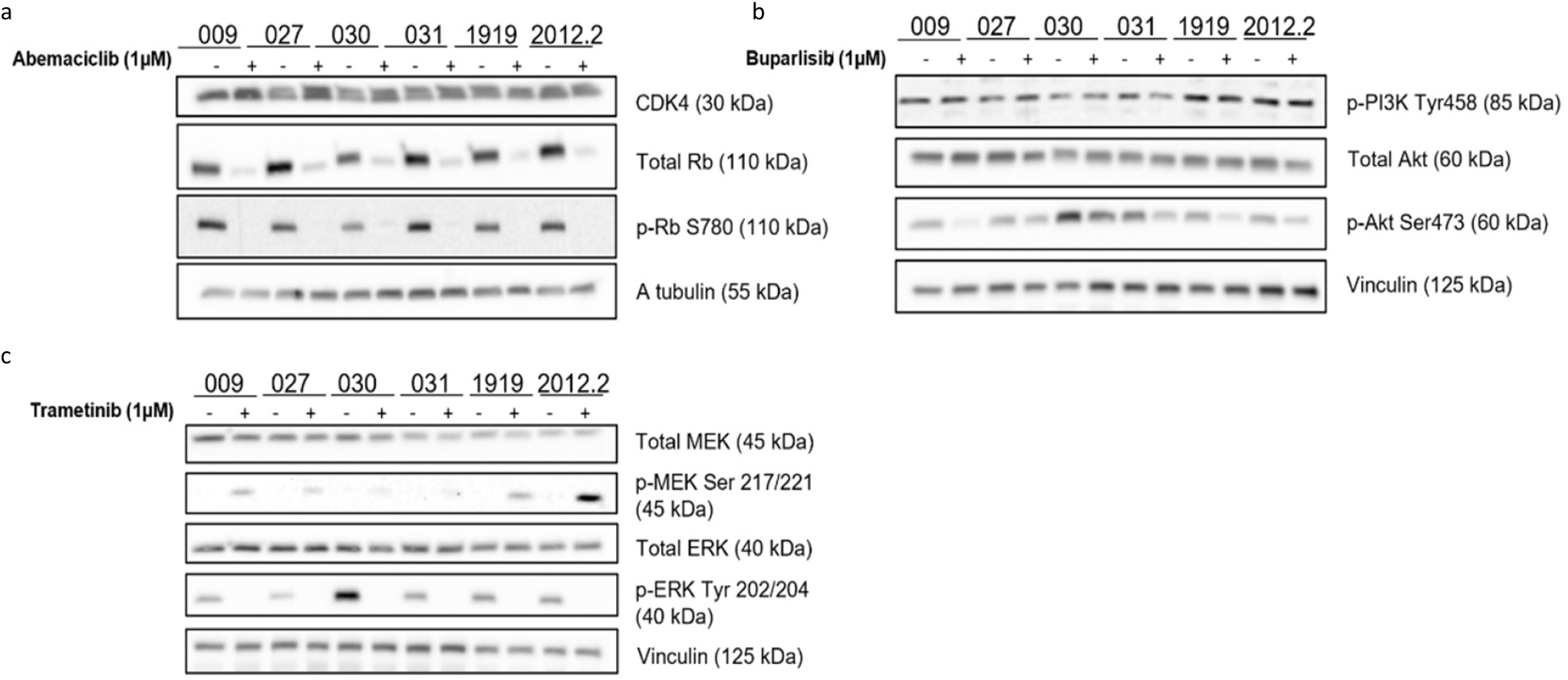
Western blot analysis of on-target efficacy of selected SMIs (1 µM, 24 hours) in PGOs. (a) Abemaciclib treatment reduced total Rb and phospho-Rb (Ser780), consistent with inhibition of the CDK4/6–Rb pathway. α-Tubulin was used as a loading control. (b) Buparlisib suppressed PI3K/AKT pathway activity, reflected by decreased phospho-AKT (Ser473) levels. Vinculin was used as a loading control. (c) Trametinib inhibited downstream phospho-ERK (Tyr202/204) expression and was associated with increased phospho-MEK (Ser217/221), consistent with MEK inhibition and upstream feedback activation. Vinculin was used as a loading control. DMSO-treated samples (indicated with a minus) served as vehicle controls.

Sunitinib reduced downstream phosphorylation specifically in PGO 2012.2, the only PGO exhibiting PDGFR-ß activation (Figure S2). Axitinib, APR-246, capmatinib, and dabrafenib failed to suppress downstream phospho-ERK at 1µM (Figure S2). Notably, dabrafenib treatment increased phospho-ERK levels in all PGOs compared to untreated controls (Figure S2). The concentrations of SMIs used for the Western blot analysis were non-cytotoxic and did not induce substantial cell death after 24 hours of treatment (Figure S2)

### 3.3 Combinatory inhibition of the MAPK pathway

Drug combinations can counteract compensatory signalling circuits activated by tumour cells exposed to single drugs. One common approach to strategies SMI drug combinations is to simultaneously inhibit multiple components of the same pathway. We sought to inhibit the MAPK and downstream pathways to prevent compensatory signalling that could lead to its reactivation. Given that EGFR signalling is one of the most prominent drivers of cellular proliferation in glioblastoma, we evaluated the effects of afatinib, trametinib, and abemaciclib as monotherapies and in combination in two molecularly distinct PGOs. PGO 1919 is characterized by pronounced MDM2 amplification, TP53 wild-type status, and a SLC16A7-CDK4 gene fusion transcript, resulting in a non-functional fusion protein. In contrast, PGO 030 harbours a potentially deleterious ERBB2 missense mutation together with CDK4 amplification and low EGFR CNV (Supplementary Table 2 and 3).

Afatinib treatment reduced pEGFR in both PGOs (Figure 2B) but only pERK expression in PGO 1919, showing a dependency of pERK activation via EGFR in this PGO. Suppression of pERK was more prominent in samples treated with trametinib in both PGOs (Figure 3A). Similarly, abemaciclib suppressed phospho-Rb activity in both PGOs (Figure 3A).

**Fig 3.**
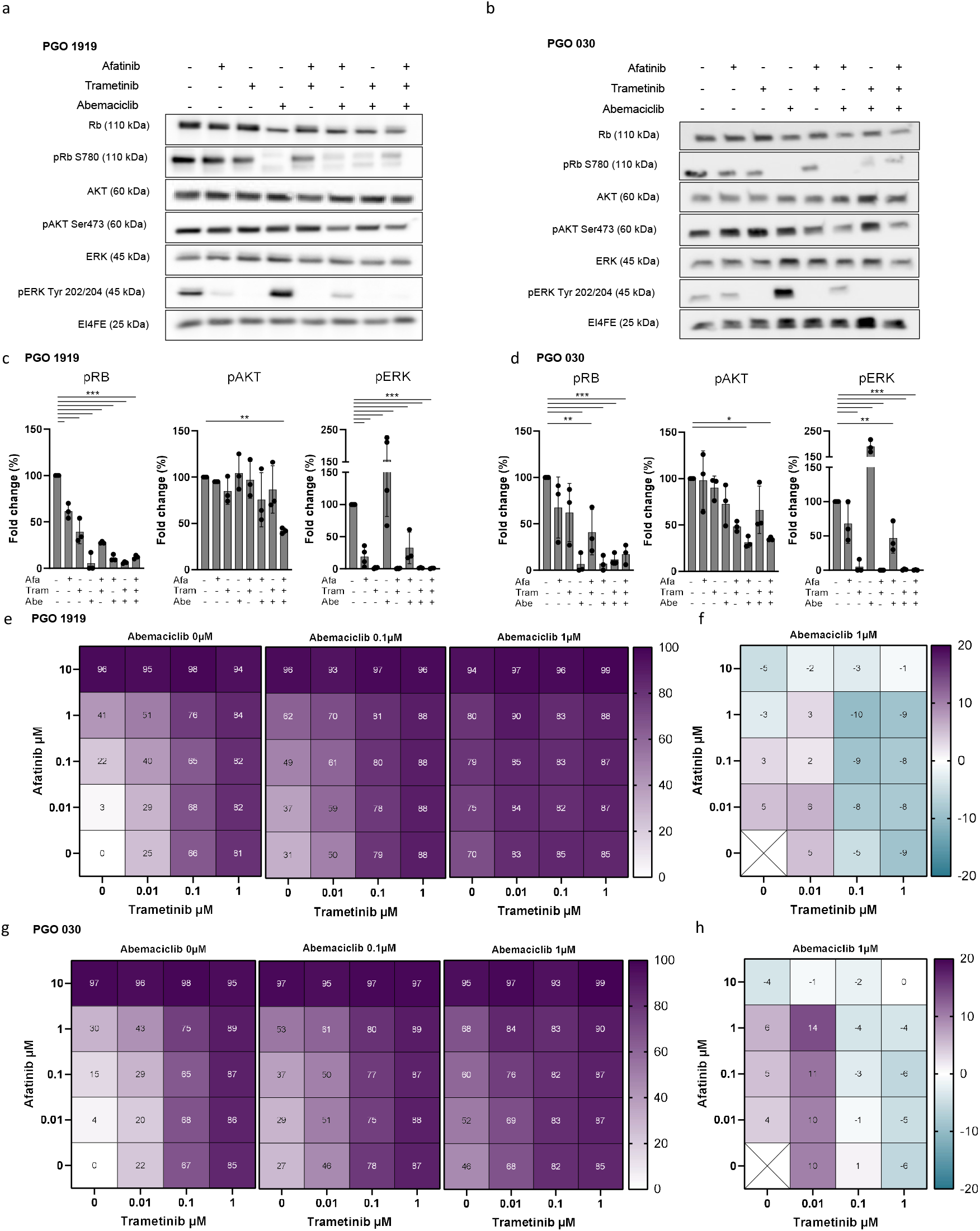
Combinatory inhibition of the MAPK pathway in PGO 1919 and PGO 030 . (a, b) PGO 1919 and PGO 030 Western blot analysis of total Rb (Rb) and phospho-Rb (pRb S780), total Akt (AKT) and phospho AKT (pAKT Ser473), total ERK and phospho ERK (pERK Tyr202/204). EI4FE and β-actin have been used as loading control. Vertically the absence or presence of SMIs was indicated by plus and minus signs e.g. with afatinib (EGFRi), trametinib (MEKi), and abemaciclib (CDK4i) 1µM, (c, d) Quantification of Western blots phosphorylated proteins, n=3 biological repeats. (e) Cell viability assay with Alamar blue showing combination treatment with afatinib (EGFRi), trametinib (MEKi), and abemaciclib (CDK4i) in PGO 1919, n=3 biological repeats (g) Cell viability assay with Alamar blue showing combination treatment with afatinib (EGFRi), trametinib (MEKi), and abemaciclib (CDK4i) in PGO 030, n=3 biological repeats (f, h) Representative Synergyfinder Bliss-based synergy scores displayed for the combination treatment with afatinib (EGFRi), trametinib (MEKi), and abemaciclib (CDK4i) in 1919 and PGO 030 respectively. Statistics were performed via ANOVA analysis. P-value is *<0.05, **<0.01, ***<0.001, ****<0.0001.

We observed that treatment with abemaciclib led to an upregulation of phospho-ERK in both PGO 1919 and 030. Importantly, this hyperphosphorylation of ERK could be blocked, with trametinib or afatinib cotreatment (Figure 3A-D). Additionally, phospho-Rb was not affected following treatment with either afatinib or trametinib, while phospho-ERK was mostly inhibited (Figure 3A-D). More significant reductions in phospho-ERK and phospho-Rb activity were observed when afatinib or trametinib were combined with abemaciclib, indicating potential synergy between EGFR or MEK inhibition and CDK4 inhibition (Figure 3A-D). To investigate this further, we conducted cell viability assays testing different combinations of the three inhibitors in PGO 1919 and 030 (Figure 3E and G). Afatinib treatment resulted in a dose-dependent reduction in cell viability, achieving a maximum inhibition of 96% at 10 µM. Similarly, trametinib induced a dose-dependent decrease, with a peak effect of 81% at 10 µM. Abemaciclib exhibited noticeable cytotoxicity, starting at 0.1 µM, with a maximum effect of 70% at 1 µM (Figures 3E and G). Combination treatment further reduced cell viability in a dose-dependent manner (Figure 3E and G).

Next, we used Synergy finder to calculate combinatorial synergy scores to determine combination treatment efficacy. In PGO 1919 and 030, the combination of afatinib with trametinib, as well as afatinib with abemaciclib, did not demonstrate synergetic interaction (synergy score < 10) (Figure 3F and H). In PGO 1919, trametinib combined with abemaciclib also lacked synergy but showed an additive effect at low concentrations, whereas, in PGO 030, a combination of 0.01 µM trametinib and 1 µM abemaciclib showed synergistic activity, with a synergy score of 10 (Figure 3F and H). This synergistic effect in PGO 030 was further enhanced by adding afatinib, resulting in synergy scores up to 10 and 14. However, the triple combination did not show a synergistic effect in PGO 1919 (Figure 3F).

### 3.4 Changes in the phospho-kinome

To more rationally strategize our combination treatment, we explored the phospho-kinome responses in PGO, via a phosphokinase array analysis on PGO 009 and PGO 031 following treatment with the three most effective inhibitors, i.e. abemaciclib, trametinib and buparlisib (Figure 4A). PGO 009 and PGO 031 were selected based on their driver mutations: IRS1 , PIK3CA/PIK3R1 and TP53, respectively (Figure S2 and 3).

**Fig 4.**
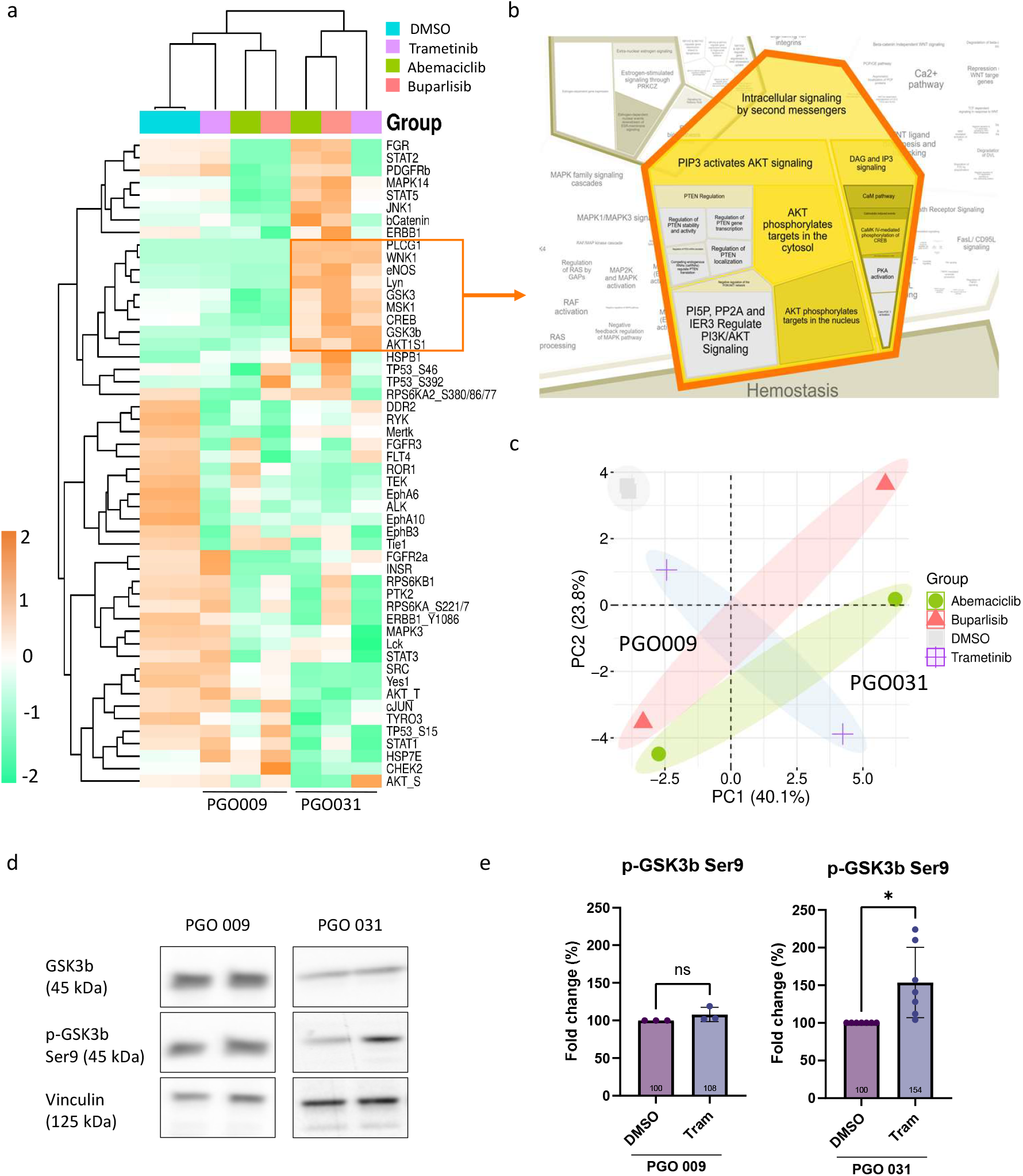
Phosphokinase array analysis of PGO responses to SMIs highlights GSK3/AKT pathway as indicator of therapeutic responsiveness. (a) Hierarchical clustering heatmap of the proteomics data was generated by SR-plot using Euclidean distance. Blue is DMSO, purple is trametinib, green is abemaciclib and red is buparlisib (b) Cluster #2 proteins presented enrichment for AKT/GSK3 pathway signalling in Reactome. (c) Principal Component Analysis (PCA) using SR-plot to determine samples and SMI response clustering using protein array proteomics analysis data. (d) Western blot for validation of total and phosphorylated GSK3β in PGO 009 and PGO 031 after 24h treatment with 1µM trametinib. Vinculin was used as a protein loading reference. (e) Densiometric quantification of western blot analysis PGO 009 n=3 and PGO 031 n=7. P-value is *<0.05.

Principal Component Analysis (PCA) demonstrates the independent clustering of both PGOs, with additional sub-clustering for the individual treatments (Figure 4C). Consistently, Euclidean hierarchical clustering separates PGOs treated with SMIs into five distinct protein clusters, within which both PGOs display differential responses to SMIs (Figure 4A). Notably, cluster #2 shows divergent signal transduction enriched for constitutive AKT/GSK3/CREB pathway activity (Figure 4A-B and Table S5). This pathway was strongly upregulated in PGO 031, but downregulated in PGO 009, underscoring pronounced differences in SMI responsiveness between the two models. The strongest difference was observed for Glycogen synthase kinase-3β (GSK-3β) phosphorylation (Ser9), which was activated in PGO 031, but not in PGO 009 (Figure 4A and D-E). Given the availability of GSK-3β inhibitors that are currently under clinical evaluation, we focused subsequent experiments on GSK-3β^Ser9^ as a potential compensatory mechanism driving resistance to these SMIs.

### 3.5 Combinatorial therapies with GSK-3ß inhibition

Phosphorylation of Serine 9 on GSK-3β (GSK-3β^ser9^), inhibits its kinase activity. Therefore, we performed cell viability assays to assess whether CHIR99021 can mimic the SMI-induced Ser9 phosphorylation, resulting in drug resistance. To confirm CHIR99021 on-target activity, we treated PGOs with 1 µM CHIR99021 for 24 hours, which led to increased inhibitory phosphorylation of GSK-3β^ser9^ compared to vehicle control in all PGOs (Figure S6). Three PGOs with different GSK-3β phosphorylation profiles were selected. In PGO 031, buparlisib and trametinib increased inhibitory phosphorylation of GSK-3β^ser9^, suggesting that GSK-3β inactivation may contribute to SMI resistance in this model. In contrast, phosphorylation levels remained unchanged in PGO 009 and PGO 027. Notably, abemaciclib reduced GSK-3β^Ser9^ across all PGOs (Figure 4D andS3).

CHIR99021 monotherapy reduced cell viability only at 10 µM across all PGOs. Combining CHIR99021 with buparlisib, abemaciclib, or trametinib further decreased cell viability in a dose-dependent manner (Figure 5A and S4). To assess potential drug interactions, Bliss synergy scores were calculated. Notably, a synergistic effect was observed in PGO 009 and PGO 027 when 0.1 µM trametinib was combined with 1 µM CHIR99021 (Figure 5B andS5). This synergistic interaction was absent in PGO 031. Optimal drug concentrations for combination treatments and synergy varied between PGOs, reflecting their distinct response profiles (Figure 5B and S5).

**Fig 5.**
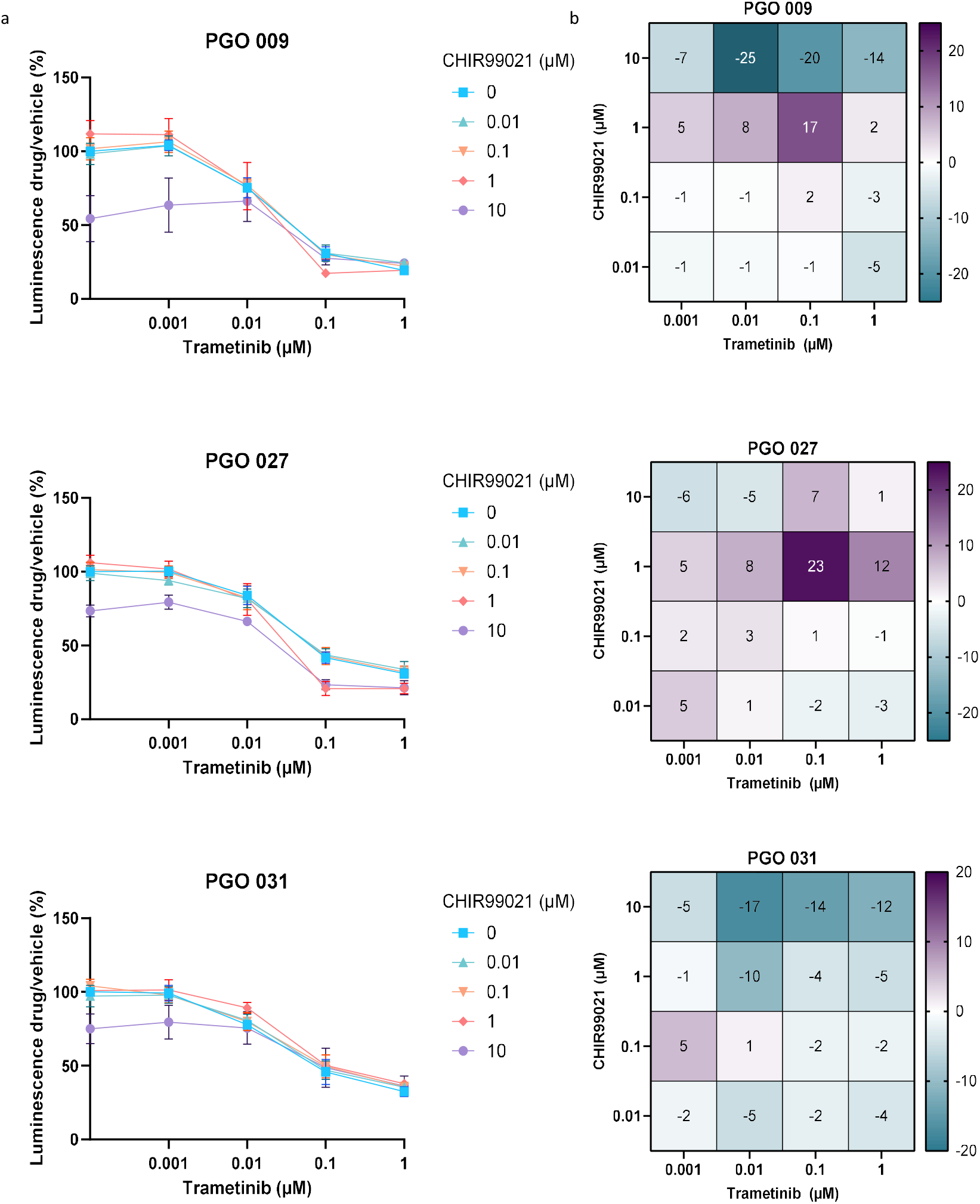
The effect of GSK-3ß inhibition in PGOs and GB patients. (a) CelltiterGlo assay was used to determine cell viability of combinatorial treatment with CHIR99021 and trametinib in PGO 009, PGO 027, and PGO 031 (n = 3 biological repeats). Data is presented as mean +/-SD (b) Synergy scores of combinatorial treatments with CHIR99021 and trametinib in PGO 009, PGO 027, and PGO 031 (n = 3 biological repeats).

### 3.6 Long term viability assays with GSK-3β inhibition

To assess the effects of trametinib and CHIR99021 combination therapy after prolonged exposure, PGO 031 was treated at multiple time points using low dosages of both drugs over 14 days (Figure 6A). Trametinib monotherapy caused a clear dose-dependent reduction in viable cell numbers, whereas CHIR99021 alone had minimal impact on viability but promoted organoid formation and increased 3D structural complexity (Figure 6B and S7A). Notably, co-treatment of CHIR99021 reversed the effects of trametinib in a CHIR99021 dose-dependent manner, indicating a clear antagonistic interaction between MEK- and GSK3β inhibition in PGO 031 (Figure 6B). Although the number of PI-negative cells per sample dropped dose-dependently with trametinib, the proportion of PI-negative cells remained similar until 12.5 nM trametinib treatment, suggesting a cytostatic rather than cytotoxic effect of the drug on low dosage (Figure 6C-F). CHIR99021 combination treatment did not alter this cytotoxicity in co-treated samples. However, similar antagonistic effects of trametinib and CHIR99021 were not seen in all GB models. In U87 GB, combined treatment with trametinib and CHIR99021 led to a more pronounced decrease in cell viability than single agent, indicating a synergistic interaction between MEK and GSK3β inhibition. Quantification using both crystal violet staining and metabolic viability assays revealed consistent trends, with Bliss synergy scores exceeding 10 (Figure S7C-F). In contrast, the U1242 GB cell line, using both crystal violet staining and metabolic viability assays, displayed clear sensitivity to CHIR99021 monotherapy, with no added benefit from the addition of trametinib. Synergy-scores displayed either additive or antagonistic effects, highlighting a clear cell model– dependent response (Figure S8).

**Fig 6.**
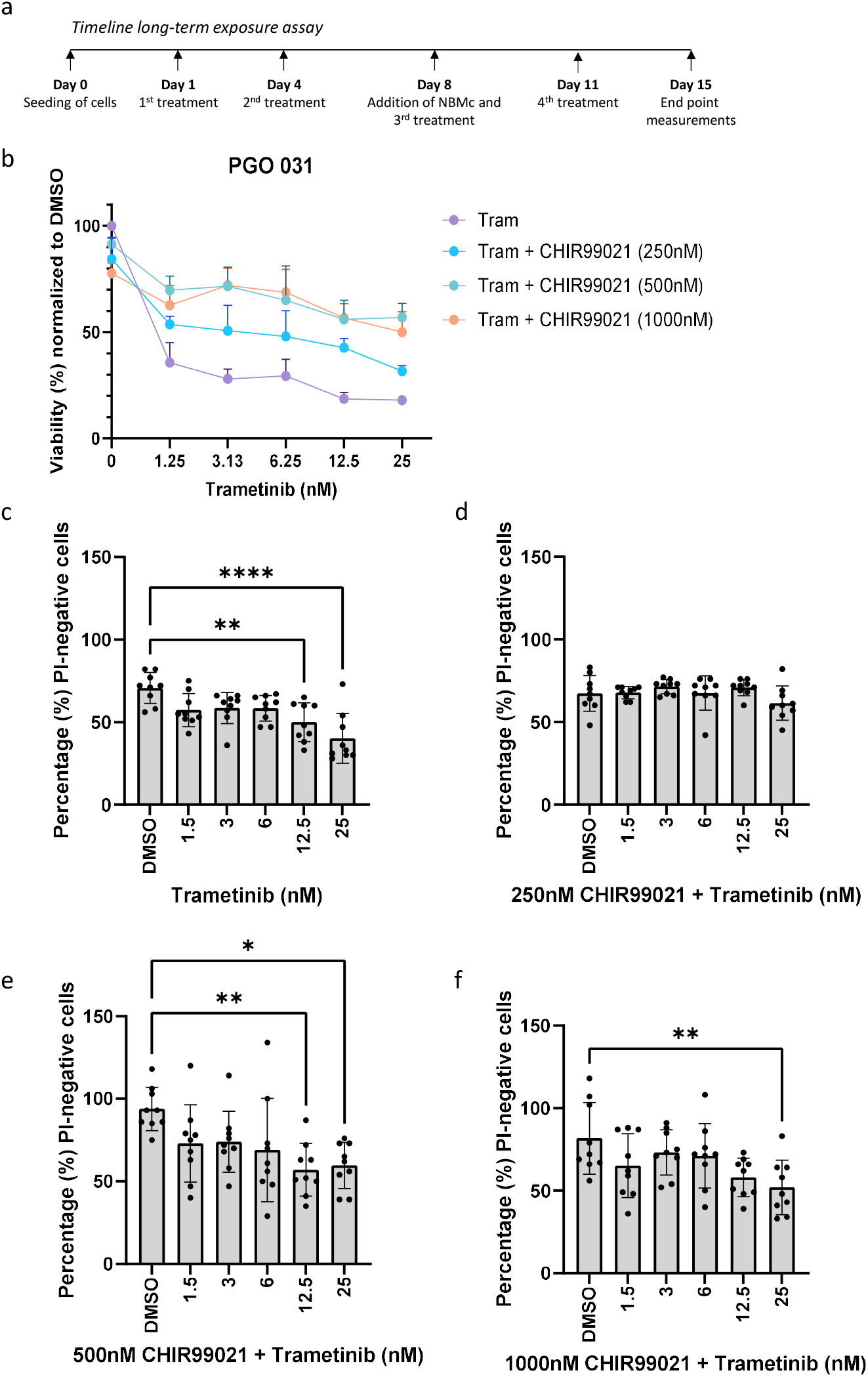
Long-term effects of GSK3ß inhibition in combination with trametinib in PGO 031. (a) Schematic overview of the experimental timeline for the long-term exposure assays. (b) Cell survival analysis, presenting the effectiveness of trametinib combined with CHIR99021, measured by flow cytometry propidium iodide (PI) staining. The average number of viable cells per sample (n=3 biological repeats), were normalized to their respective DMSO vehicle control, average ± SD (c - f) Percentage cell viability after trametinib, or the individual combinations, as measured by flow cytometry PI-staining per sample (n=3 biological repeats, with three technical repeats). Statistics were performed via ANOVA analysis. P-value is *<0.05, **<0.01, ***<0.001, ****<0.0001.

## 4. Discussion

We evaluated a focused panel of genetically characterised patient-derived GB organoids (PGOs) to interrogate kinase pathway dependencies and adaptive responses to clinically approved SMIs. Across PGO models, on-target pathway inhibition was achieved, yet the magnitude of pathway suppression and growth inhibitory effects differed substantially. Importantly, single-agent treatment induced compensatory signalling that was PGO-specific, enabling the identification of actionable bypass nodes and rational combinations to mitigate adaptive resistance Across this panel, inhibitors targeting downstream signalling effectors (abemaciclib, trametinib, buparlisib) generally produced more consistent pathway suppression and viability reduction than upstream RTK inhibitors (i.e. afatinib, capmatinib). This is consistent with the concept that downstream blockade can constrain convergent oncogenic inputs and reduce pathway redundancy relative to RTK-directed inhibition, which is more vulnerable to receptor-level bypass (33, 34). Nevertheless, the relationship between mutational profiles and drug response was modest, underscoring that single alterations are insufficient to predict dependency in these models, where multiple co-occurring mutations and cell-state programs rewire oncogenic signalling. Targeted kinase inhibitors often produce rapid but transient responses because signalling networks adapt through release of negative feedback regulation, and compensatory pathway activation, restoring downstream signalling. In addition, GB cell populations are heterogeneous and pre-existing less responsive cell clones might enable survival and outgrowth of resistant cell states. In EGFR-mutant NSCLC, acquired resistance to EGFR TKIs is a dynamic process driven by secondary EGFR mutations (e.g., T790M), MET amplification, and phenotypic switching, with occasional reversion of drug sensitivity (35). Clinically, the benefit of rational combinations is exemplified by EGFR plus MET inhibition in relapsed NSCLC (34, 36). These findings highlight the importance of functional pathway mapping upfront to complement genomic profiling when designing stratification strategies and combination trials such as the Drug Rediscovery Protocol Study (DRUP) (37, 38).

Importantly, we observed effective target inhibition with SMIs across all PGO models, warranting further research to identify optimal treatment schedules and combinations of these inhibitors for GB patients. A phase 0 clinical trials in GB has demonstrated that the CDK4/6 inhibitor ribociclib achieves concentrations up to fivefold the in vitro IC50 in both the cerebrospinal fluid and the tumor, suggesting that inadequate CNS/tumor penetration is not the sole barrier for limited treatment efficacy (39). Instead, adaptive rewiring and intratumoral heterogeneity can rapidly reduce target dependency (6, 40, 41). For instance, the inhibition of CDK4/6 activates c-MET and Tropomyosin receptor Kinase A-B (TrkA-B) pathways via Nuclear factor kappa-light-chain-enhancer of activated B cells (NF-kB) signalling in GB (25, 41). Furthermore, CDK4/6 inhibition led to compensatory activation of the MAPK- and PI3K/AKT pathway, indicating the presence of a feedback loop that reinforces cell survival (39, 40, 42). This observation is consistent with our finding of upregulated p-ERK levels after treatment with abemaciclib. Interestingly, abemaciclib treatment also led to a reduction in total Rb protein levels across all PGOs tested. As abemaciclib maintains Rb protein in a hypophosphorylated state bound to E2F, the availability of free E2F is likely diminished. Given that the *RB1* promoter contains an E2F-binding site, reduced E2F availability may attenuate transcriptional activation of the *RB1* gene itself, thereby contributing to the decreased total Rb protein levels observed (43).

Due to pathway rewiring, patient selection based on bulk resected tumor profiles is unlikely to reliably identify actionable dependencies that translate into durable responses. Instead, a mechanistic understanding of treatment-induced kinome rewiring is required to design combinations that prevent or overcome adaptive resistance and suppress escape. Blocking at multiple levels within the same signalling cascade may achieve more sustained and complete inhibition of the pathway, preventing adaptive feedback mechanisms that lead to drug resistance (44-48). Clinical studies have demonstrated the efficacy of this strategy for instance, the combination of dabrafenib and trametinib has improved overall survival in melanoma patients with *BRAF V600E* or *V600K* mutations (49). In patients with BRAF wild type, treatment with dabrafenib has shown to induce ERK phosphorylation, al observed here in GB. Interestingly, we observed trametinib-induced hyperphosphorylation of MEK at Ser217/221 in all PGO models, suggestive for a reactivation of the MAPK pathway through an unknown upstream cascade. Despite compensatory pathway activation, trametinib remained effective in suppressing downstream ERK signalling, likely by inhibiting MEK Ser217 phosphorylation, thereby preventing the dual phosphorylation of MEK1/2 required for ERK activation (50). Combining trametinib with an SMI targeting this compensatory signal rerouting could potentially result in an effective synergistic drug combination and more potent cell killing.

To further investigate the potential of vertical inhibition strategies in GB, we examined the combination of afatinib, trametinib, and abemaciclib in two PGOs. Trametinib induced a more pronounced reduction in pERK levels in both PGOs compared with upstream RTK inhibition. Afatinib reduced pERK selectively, but not completely, in PGO 1919, indicating differential dependency on upstream receptor signalling between the PGO models. Similar results were obtained by a phase II clinical trial in GB, in which patients were treated with the EGFR-inhibitor gefitinib. Initially high concentrations of the drug were found in tumor tissue, and on-target inhibition of EGFR was obtained, but the drug failed to subsequently downregulate the MAPK pathway (51). In addition, the effectiveness of gefitinib did not correlate with EGFR mutational status, indicating that the signalling network involving other ERBB family members and TKRs such as MET and PDGFR is robust, owing to its modular organization and redundancy in regulatory circuits.

Neither EGFR nor MEK inhibition alone was sufficient to fully induce cell cycle arrest, supporting the rationale for combining these agents with CDK4/6 inhibition to achieve enhanced anti-proliferative effects. Notably, CDK4/6 blockade induced ERK hyperphosphorylation, which was effectively counteracted by co-treatment with a MEK inhibitor. In PGO 1919, afatinib more effectively attenuated abemaciclib-induced ERK hyperphosphorylation. These observations suggest that CDK4/6 inhibition reactivates the MAPK pathway, through upstream RTK activation. Similar feedback mechanisms have been described in head and neck squamous cell carcinoma, where CDK4/6 inhibition induced ERK hyperphosphorylation that was reversed by trametinib (46), and in ER-positive breast cancer, where EGFR and HER2 mediated ERK reactivation following CDK4/6 blockade (52). Importantly, combined MEK and CDK4/6 inhibition was synergistic in PGO 030 but not in PGO 1919. This likely reflects the greater dependency of PGO 1919 to EGFR or MEK inhibitor monotherapy, which already substantially induces anti-proliferative effects.

Subsequently, we used phosphokinase arrays to map signalling responses to abemaciclib, buparlisib, and trametinib, revealing these SMIs rewire the AKT/GSK3/CREB axis in a PGO-dependent manner, with GSK3β showing the greatest divergence. Notably, trametinib–GSK3β inhibitor antagonism may result from feedback activation of WNT/β-catenin signalling. GSK3β is a central negative regulator of WNT, as it phosphorylates β-catenin and promotes its degradation (53, 54). Inhibition of GSK3β, either through AKT-mediated Ser9 phosphorylation or through pharmacological blockade, prevents its kinase activity, stabilizes β-catenin, and activates the WNT pathway (55). This activation of WNT signalling is well known to enhance stemness and therapy resistance in GB (56). Consistent with this, previous reports have shown that MEK inhibition can trigger adaptive signalling through PI3K/AKT, leading to inhibitory phosphorylation of GSK3β at Ser9, nuclear accumulation of β-catenin, and upregulation of stemness programs (57). In our study, trametinib induced AKT activation and Ser9 phosphorylation of GSK3β specifically in PGO 031, providing a mechanistic rationale for the observed antagonism when combined with a GSK3β inhibitor. This is supported by the observation that GSK3β inhibition promoted the formation of larger and more compact 3D cell clusters in GB PGOs, which may reflect enhanced stemness and self-renewal capacity.

While the specific drug combinations explored in this study are novel, one previous study has investigated the combination of CHIR99021 and trametinib. This dual treatment more effectively reduced the growth of orthotopic human PDAC xenografts in mice compared to monotherapy (58). Beyond this, other GSK3β inhibitor-based combinations have shown promise in various cancer types. For instance, combined GSK3β and poly ADP-ribose polymerase (PARP) inhibition demonstrated strong synergy in a genetically diverse panel of colorectal cancer cell lines (59). In another study, the GSK3 inhibitor LY2090314 synergized with the anaplastic lymphoma kinase (ALK) inhibitor lorlatinib in intermediate-resistant ALK-positive NSCLC cells (60). These findings highlight a context-dependent role of GSK3β in GB. Further studies dissecting the interplay between MEK, PI3K/AKT, and WNT signalling in GB will be critical to identify subgroups where combined pathway inhibition is therapeutically beneficial versus detrimental.

This work has several limitations. Drug concentrations used in mechanistic Western blot experiments may exceed clinically achievable exposures, potentially increasing off-target effects and limiting translational relevance. We note that the, concentrations used in cell viability assays however are within clinically relevant ranges. Clinical PK studies of the MEKi avutometinib, the EGFRi gefitinib, and the CDK4/6i ribociclib report unbound drug concentrations in GB tissue that exceed the IC50 values described in this study (39, 51, 61). Further pharmacogenetic and pharmacokinetic studies of SMIs in the GB brain compartment are needed to define optimal dosing strategies for combination treatments. Additionally, the toxicity of combining three SMIs could be significant and potential clinically relevant combination regimens should be carefully tested in a phase I clinical trials (62). However, the primary objective here was to uncover compensatory signalling networks and nominate rational combinations, not to define clinical dosing. A second limitation is cohort size, which precludes robust genotype– response conclusions and restricts generalization. Future studies should scale functional profiling across additional models and integrate orthogonal readouts (e.g. phosphoproteomics and single-cell analyses) to refine predictive signatures of adaptive rewiring and to prioritize combinations most likely to sustain pathway suppression at clinically feasible exposures.

Current patient stratification strategies often focus on single-driver mutations and rely on limited temporal sampling to guide treatment decisions, providing only limited benefit. Our findings provide molecular insights that support the conclusion that stratifying patients by single-driver mutations is insufficient, as resistance, heterogeneity, and pathway activation are driven by the broader mutational context. Achieving durable benefit from SMI therapies will require precise patient stratification, temporal monitoring and rational combination of SMIs to suppress compensatory signalling and improve clinical responses (35, 63).

In summary, our data support a framework in which GB resistance to targeted therapy is driven in part by rapid, context-specific kinome adaptation. Mechanistic dissection of these compensatory responses can reveal actionable bypass nodes and guide rational combinations, with the goal of improving the durability of targeted interventions in GB.

## Supporting information

Suppkementary Figures

## Data accessibility

All data supporting the findings of this study are included in the main text and *Supplementary Data*. Additional raw data files generated and analysed during the current study are available from the corresponding author upon reasonable request.

## Acknowledgements

We thank the patients who donated material and the surgeons, assistants, and nurse practitioners who helped acquiring the tumor material. We thank Maikel Verduin, Bartosz Olaszkiewicz and Lisa Duijcx (Maastricht University) for their help during the project.

## Author Contribution

Conceptualization - LH, MvH, AH and MV. Software – KK. Investigation - LH, MvH and JP. Resources – SR. Writing of the original draft – LH and MvH. Writing, reviewing and editing – all authors contributed. Visualization – MvH. Supervision – AH and MV.

## Statements and declarations

This scientific publication was supported by funding KWF Kankerbestrijding (INTO-PROT grant #12092) and Stichting STOPHersentumoren. The authors declare that they have no conflicts of interest or competing financial interests. The authors used ChatGPT (OpenAI, GPT-5.5) to assist with language editing and improvement of manuscript readability; all scientific content, interpretation, and conclusions were reviewed and verified by the authors.

