## Supplementary material for "Adaptive Kinase Signalling Enables Escape from Small Molecule Inhibition in Glioblastoma": Suppkementary Figures

**Supplementary methods**

*Supplementary Table 1: Used primary and secondary antibodies for Western blot*

| **Antibody** | **Company** | **Product number** |
|---|---|---|
| PDGFR-β | Cell Signalling Technology | 3169 |
| phospho-PDGFR-β Tyr1009 | Cell Signalling Technology | 3124 |
| PI3K | Cell Signalling Technology | 4257 |
| phospho-PI3K Tyr458/Tyr199 | Cell Signalling Technology | 4228 |
| EGFR | Cell Signalling Technology | 4267 |
| phospho-EGFR Tyr1068 | Cell Signalling Technology | 2234 |
| VEGFR-2 | Cell Signalling Technology | 2479 |
| phospho-VEGFR-2 Tyr1175 | Cell Signalling Technology | 2478 |
| c-KIT | Cell Signalling Technology | 3780 |
| phospho-c-KIT Tyr703 | Cell Signalling Technology | 3073 |
| c-MET | Cell Signalling Technology | 8198 |
| phospho-c-MET Tyr1234/1235 | Cell Signalling Technology | 3077 |
| p53 | Santa Cruz | SC-476981043 |
| phospho-p53 S15 | R&D systems | AF1043 |
| p21 | BD-Pharmingen | 556430 |
| Akt | Cell Signalling Technology | 4691 |
| phospho-Akt | Cell Signalling Technology | 4060 |
| ERK | Cell Signalling Technology | 4695 |
| phospho-ERK | Cell Signalling Technology | 4370 |
| MEK | Cell Signalling Technology | 4694 |
| phospho-MEK S217/221 | Cell Signalling Technology | 9121 |
| Rb | Cell Signalling Technology | 9309 |
| phospho-Rb Ser780 | Cell Signalling Technology | 9307 |
| CDK-4 | Cell Signalling Technology | 12790 |
| alpha-Tubulin | Sigma | T6199 |
| Vinculin | Sigma | V9131 |
| EI4FE | Cell Signalling Technology | 9741 |
| B-actin | Cell Signalling Technology | 4967 |
| PLC-γ1 | Cell Signalling Technology | 5690 |
| phospho-PLC-γ1 Tyr783 | Cell Signalling Technology | 2821 |
| GSK3ß | Cell Signalling Technology | 12456 |
| phospho-GSK3ß Ser9 | Cell Signalling Technology | 5558 |
| Lyn | Cell Signalling Technology | 2796 |
| phospho-Lyn Tyr397 | Cell Signalling Technology | 70926 |

| | | |
|---|---|---|
| Chk2 | Cell Signalling Technology | 2662 |
| phospho-Chk2 Thr68 | Cell Signalling Technology | 2197 |
| anti-rabbit | Cell Signalling Technology | 7074 |
| anti-mouse | Cell Signalling Technology | 7076 |

*Supplementary Table 2: complete overview of Log2 values copy number variants in PGOs*

| **Log2 values CNV-analysis** | | | | | |
|---|---|---|---|---|---|
| | **PGO009** | **PGO027** | **PGO030** | **PGO031** | **PGO1919** |
| EGFR | 1.207 | 1.364 | 0.395 | 2.750 | 1.580 |
| PDGFRA | -0.095 | 0.057 | -0.368 | 0.388 | |
| CDK4 | 4.374 | -0.021 | 4.446 | 0.171 | |
| KIT | 0.627 | 0.113 | -0.306 | 0.256 | |
| MDM2 | 2.341 | -0.509 | 6.277 | -0.466 | 6.687 |
| PTEN | 4.788 | 0.134 | 0.633 | -0.016 | |
| NF1 | 0.060 | -0.754 | 0.392 | -0.144 | |
| CDKN2C | -0.420 | -0.925 | -0.567 | -0.435 | |
| CHD5 | 0.063 | -0.168 | 0.097 | -0.015 | |
| MET | 0.388 | 0.451 | 0.711 | -0.268 | |
| CDK6 | 0.671 | 0.853 | 0.513 | -0.165 | |
| TP53 | 0.939 | 0.880 | 0.214 | -0.047 | |
| CCND2 | 0.359 | 0.136 | 0.053 | -0.928 | |
| PIK3CA | 0.138 | -0.012 | 0.517 | 0.359 | |
| RB1 | 0.021 | -0.052 | 0.374 | 0.536 | |

*Supplementary Table 3: Mutational characteristics of PGOs (33)*

| **Organoid line** | **Pathogenic mutations present (based on WES)** | **Mutated in GBM (TCGA)** | **SIFT** | **CADD** | **ENSEMBL ID** | **COSMIC ID** | **Conclusion** |
|---|---|---|---|---|---|---|---|
| **PGO 009** | *IRS1 p.Gly818Arg* | yes | 0.01 | 23 | *rs41265094* | *COSV99043455* | Predicted to affect protein functioning |
| | *EPHA2 p.Arg93Cys* | - | 0 | 32 | *rs200152244* | - | Strongly predicted to affect protein functioning |
| | *MKNK1 p.Arg310Gln* | yes | - | - | *Unknown* | - | - |
| | *PRKAG3 p.Asp485Asn* | yes | *0.06* | *21* | *rs149508864* | - | Likely benign |
| **PGO 027** | *NF1 p.Lys1423Gln* | yes | - | - | *MDB00617536* | - | Pathogenic/likely pathogenic |
| | *RASGRF1 pAsn218His* | yes | - | - | *Unknown* | - | - |
| | *TAOK1 p.Thr185Ala* | yes | - | - | *Unknown* | - | - |
| | *EphA1 p.Arg88His* | yes | *0* | *28* | *rs148653512* | - | Potential deleterious |
| | *PTEN p.Val158Gly* | yes | - | - | *Unknown* | - | - |
| | *CDK11A p.Arg327Trp* | - | - | - | *Unknown* | - | - |
| | *TRRAP p.Thr2076Arg* | - | - | - | *Unknown* | - | - |
| | *TAF1 p.Ser547Cys* | yes | - | - | *Unknown* | - | - |
| | *SKA3 p.Arg154Cys* | yes | 0.08 | 19 | *rs1256196715* | *COSV53437349* | Likely benign |
| **PGO 030** | *EphA2 p.Arg93Cys* | - | 0 | 32 | *rs200152244* | - | Likely deleterious |
| | *EphA3 p.Ala777Gly* | yes | 0.01 | 27 | *rs34437982* | - | Potential deleterious |

| | | | | | | | |
|---|---|---|---|---|---|---|---|
| | *ERBB2 p.Ala1216Asp* | yes | 0 | 23 | *rs55943169* | *COSV99059727* | Potential deleterious |
| | *IRS1 p.Gly818Arg* | yes | 0.01 | 23 | *rs41265094* | *COSV99043455* | Potential deleterious |
| | *FGFR3 p.Pro451Ser* | yes | - | - | *Unknown* | - | - |
| | *FOXO3 p.Ala140Val* | yes | 0.16 | 22 | *rs111556510* | *COSV59628700* | Likely benign |
| | *RASAL1 p.Ala93Val* | yes | 0.54 | 18 | *rs146551951* | - | Benign |
| | *MET p.Asn375Ser* | yes | 0.1 | 19 | *rs33917957* | *COSV59260365* | Benign |
| **PGO 031** | *PIK3CA p.Asn1044Lys* | *yes* | 0.04 | 22 | *rs1576949960* | *COSV55897116*<br>*COSV55875087* | Potential deleterious |
| | *PIK3R1 p.Pro125Leu* | *yes* | 0 | 26 | *rs1401537376* | - | Potential deleterious |
| | *CDK15 p.Gly75Ala* | - | 0 | 24 | *rs1336667433* | *COSV53632739* | Potential deleterious |
| | *TP53 p.Cys176Trp* | *yes* | 0 | 20 | *rs1057519980* | *COSV99377692*<br>*COSV52735850*<br>*COSV52701599*<br>*COSV52687391* | Potential deleterious |
| **1919** | *ERBB2 p.Gly201Val* | yes | 0.12 | 2 | *rs750651941* | *COSV99506529*<br>*COSV105036629* | Likely benign |
| | *JAK3 p.Arg887His* | yes | 0.06 | 23 | *rs148688786* | *COSV105940658* | Likely benign |
| | *RB1 p.Cys590* (STOP)* | yes | - | - | *rs145310579* | *COSV99031794*<br>*COSV57295755* | Loss of function |
| | *SLC16A7–CDK4 fusion* | - | - | - | - | - | Likely loss of function |
| **2012.2** | *EphA3 p.Ala777Gly* | yes | 0.01 | 27 | *rs34437982* | *COSM1232876* | Potential deleterious |

| | | | | | | | |
|---|---|---|---|---|---|---|---|
| | *ERBB2 p.Ala1216Asp* | yes | 0 | 23 | *rs55943169* | *COSV99059727* | Potential deleterious |
| | *FGFR3 p.Pro449Ser* | yes | 0 | 23 | *rs61735104* | COSV53422884 | Potential deleterious |
| | *LTK p.Arg569Ser* | - | 1 | 17 | *rs148513655* | *COSV53027475 COSV53026607 COSV107230095* | Likely benign |

*Supplementary table 4: Selected SMIs with their targets*

| **Compound** | **Supplier** | **Targets ordered by affinity (highest to lowest)** |
|---|---|---|
| Abemaciclib | Bio-Connect (1231929-97-7) | CDK4, CDK6 |
| Afatinib | Bio-Connect (850140-72-6) | EGFR, ERBB4, ERBB2 |
| APR-246/PRIMA-1MET | Bio-Connect (5291-32-7) | TP53 |
| Axitinib | Bio-Connect (319460-85-0) | VEGFR1, VEGFR2, VEGFR3, PDGFRß, c-Kit |
| Buparlisib | Bio-Connect (HY-70063) | PI3K |
| Capmatinib | Bio-Connect (HY-13404) | MET |
| Dabrafenib | Bio-Connect (HY14-660) | BRAF V600E |
| Sunitinib | Bio-Connect (557795-19-4 | PDGFRß, c-Kit, VEGFR2, PDGFRa, VEGFR1, FLT3, VEGFR3, RET, CSF-1R |
| Trametinib | Bio-Connect (HY-10999) | MEK1, MEK2 |

*Supplementary table 5 clustering analysis of SMI treated PGOs*

| **Cluster #2**<br>**Pathway name** | proteins found | pathway total | pValue | FDR | Reactions found | Reactions total |
|---|---|---|---|---|---|---|
| Constitutive Signaling by AKT1 E17K in Cancer | 4 | 29 | 5.20E-09 | 1.68E-06 | *3* | *18* |
| AKT phosphorylates targets in the cytosol | 3 | 16 | 2.16E-07 | 3.48E-05 | *2* | *9* |
| Diseases of signal transduction by growth factor receptors and second messengers | 6 | 536 | 1.07E-06 | 1.15E-04 | *44* | *516* |
| PI3K/AKT Signaling in Cancer | 4 | 132 | 2.14E-06 | 1.71E-04 | *3* | *21* |
| Intracellular signaling by second messengers | 5 | 362 | 4.04E-06 | 2.58E-04 | *7* | *116* |
| CREB phosphorylation | 2 | 9 | 2.06E-05 | 4.13E-04 | *4* | *4* |
| Transcriptional and post-translational regulation of MITF-M expression and activity | 3 | 79 | 2.53E-05 | 4.33E-04 | *5* | *37* |
| MECP2 regulates transcription factors | 2 | 10 | 2.55E-05 | 4.33E-04 | *2* | *8* |
| Axon guidance | 5 | 566 | 3.50E-05 | 5.94E-04 | *21* | *295* |

### Supplementary Figures

#### Supplementary figure 1

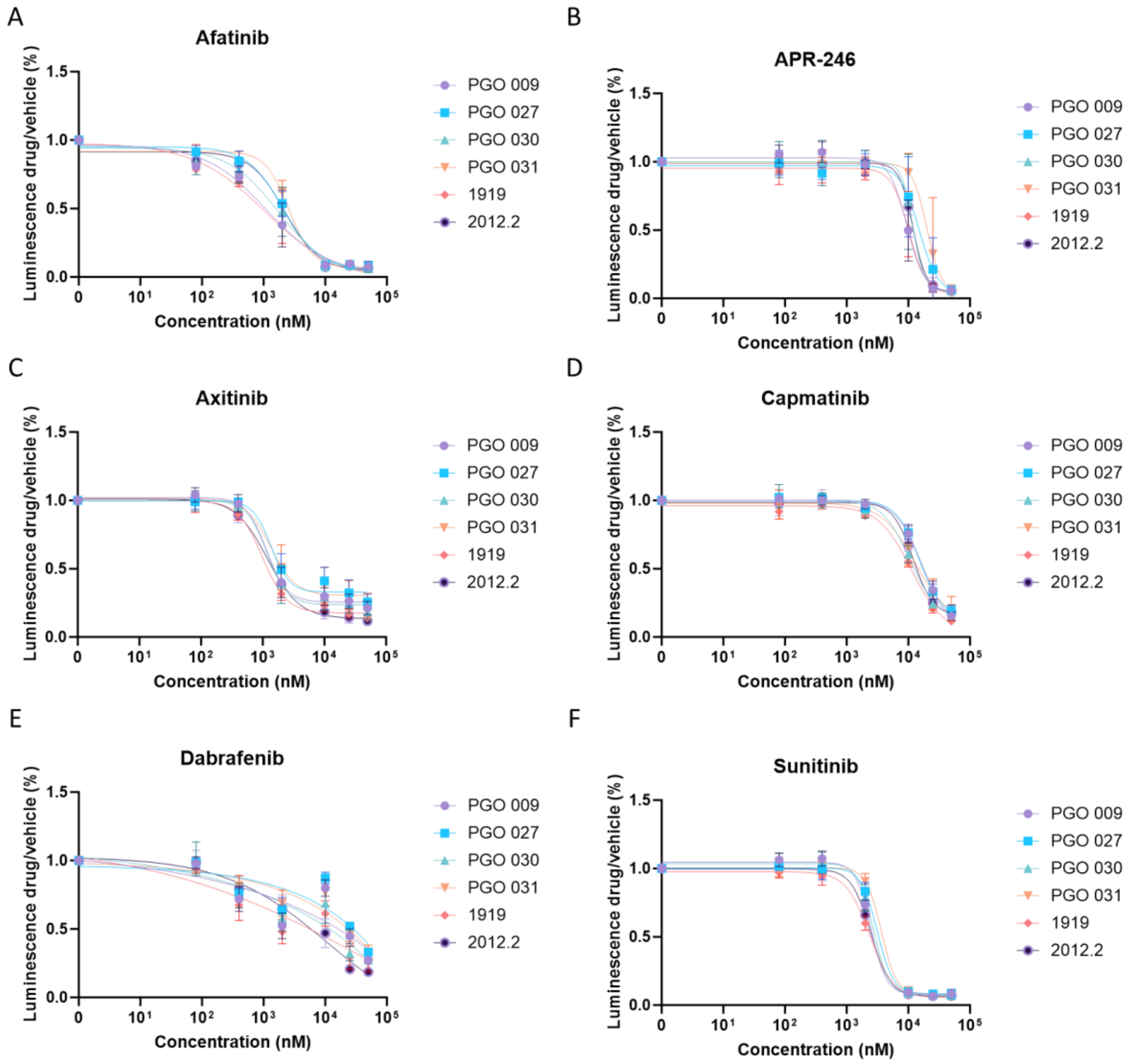

*Supplementary Figure 1: Treatment sensitivity of patient-derived glioblastoma organoids (PGOs) to selected small-molecule inhibitors (SMIs).* PGOs were treated with increasing concentrations of the indicated inhibitors for 7 days, after which cell viability was measured using the CellTiter-Glo luminescence assay. Dose–response curves are shown for (A) afatinib (EGFR inhibitor), (B) APR-246 (restores p53 function), (C) axitinib (VEGFR, PDGFRβ, and c-Kit inhibitor), (D) capmatinib (MET inhibitor), (E) dabrafenib (BRAF V600E inhibitor), and (F) sunitinib (PDGFRβ inhibitor). Data represent mean ± SD of N = 3 independent biological replicates.

**Supplementary figure 2**

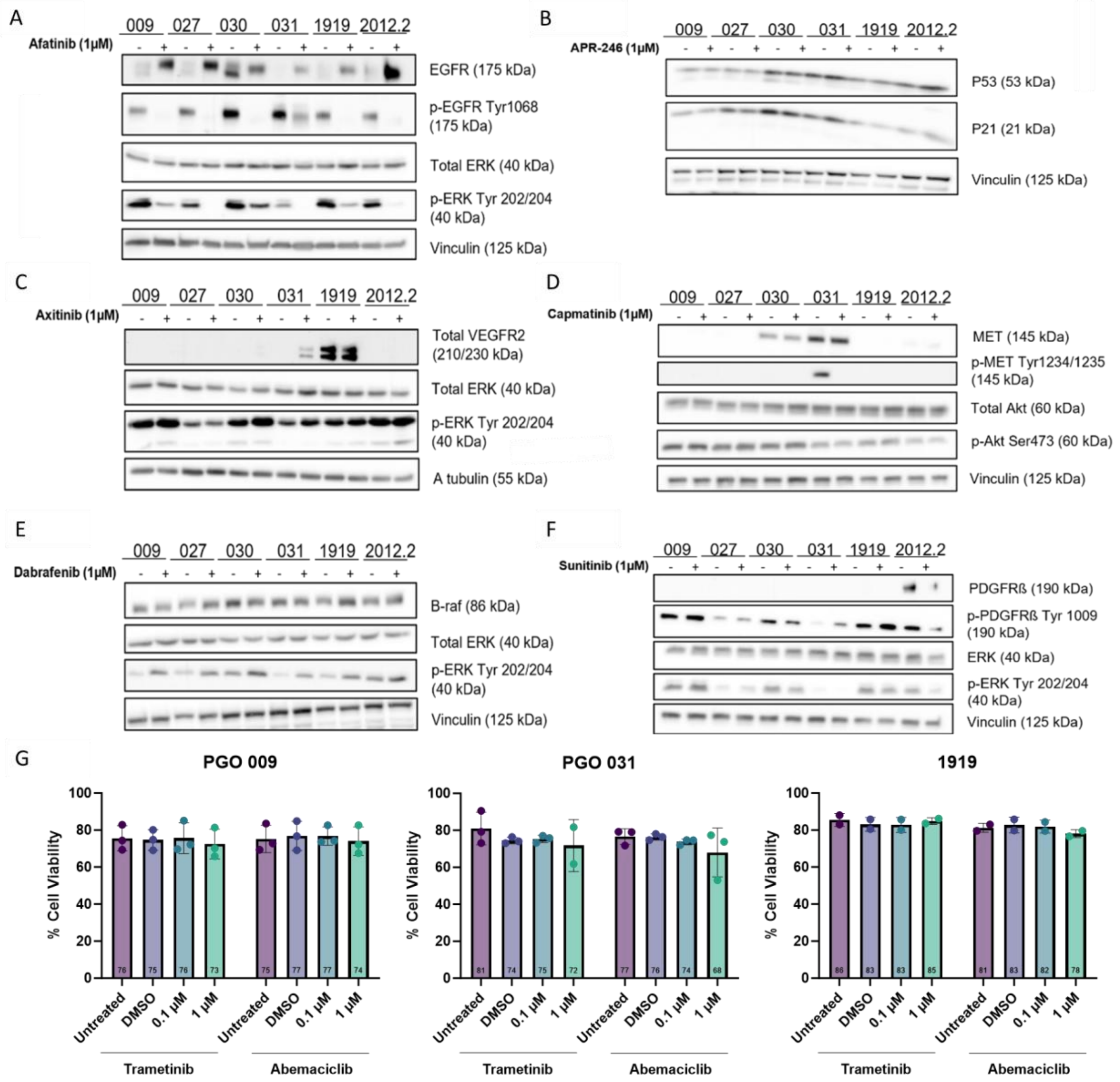

*Supplementary Figure 2: On-target efficacy of small-molecule inhibitors Western blot analysis of PGOs following treatment with the indicated inhibitors (1 µM, 24 h):* (A) afatinib, assessing EGFR, phospho-EGFR, ERK and phospho-ERK; (B) APR-246, assessing p53 (tumor protein p53) and p21 (cyclin-dependent kinase inhibitor 1); (C) axitinib, assessing VEGFR, ERK, and phospho-ERK; (D) capmatinib, assessing MET, phospho-MET, Akt, and phospho-Akt (E) dabrafenib, assessing B-Raf, ERK, and phospho-ERK; and (F) sunitinib, assessing PDGFRβ, phospho-PDGFRβ, ERK, and phospho-ERK. Vinculin was used as a loading control in all experiments. (G) Flow proprium iodide staining after 24h of treatment with trametinib and abemaciclib at two different concentrations (0.1 and 1µM). DMSO was used as a vehicle control.

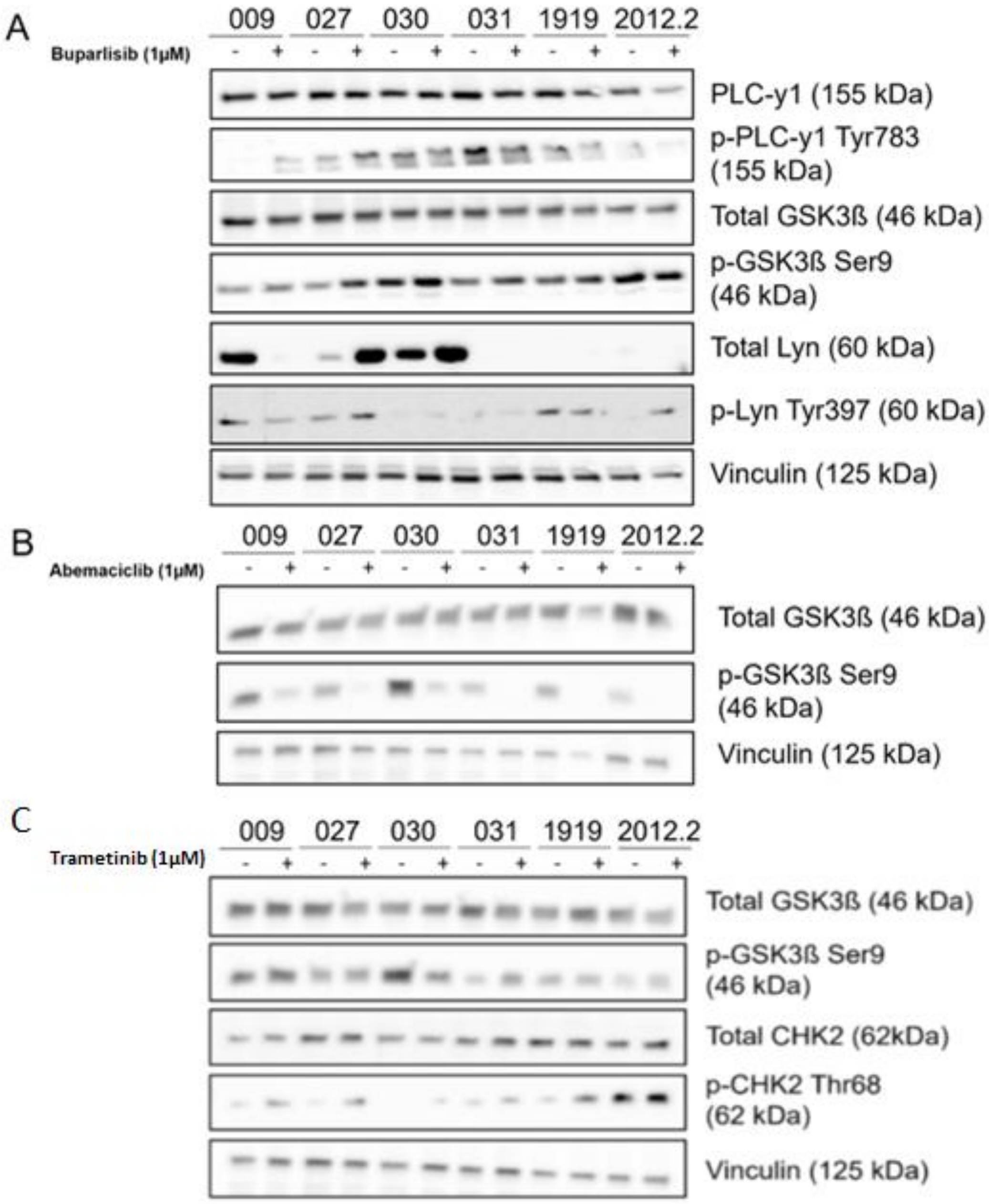

*Supplementary Figure 3: Western blot for validation of phosphokinase array in all PGOs after 24h treatment with 1µM buparlisib (A), abemaciclib (B), trametinib (C) (N = 1).* Vinculin was used as a loading control.

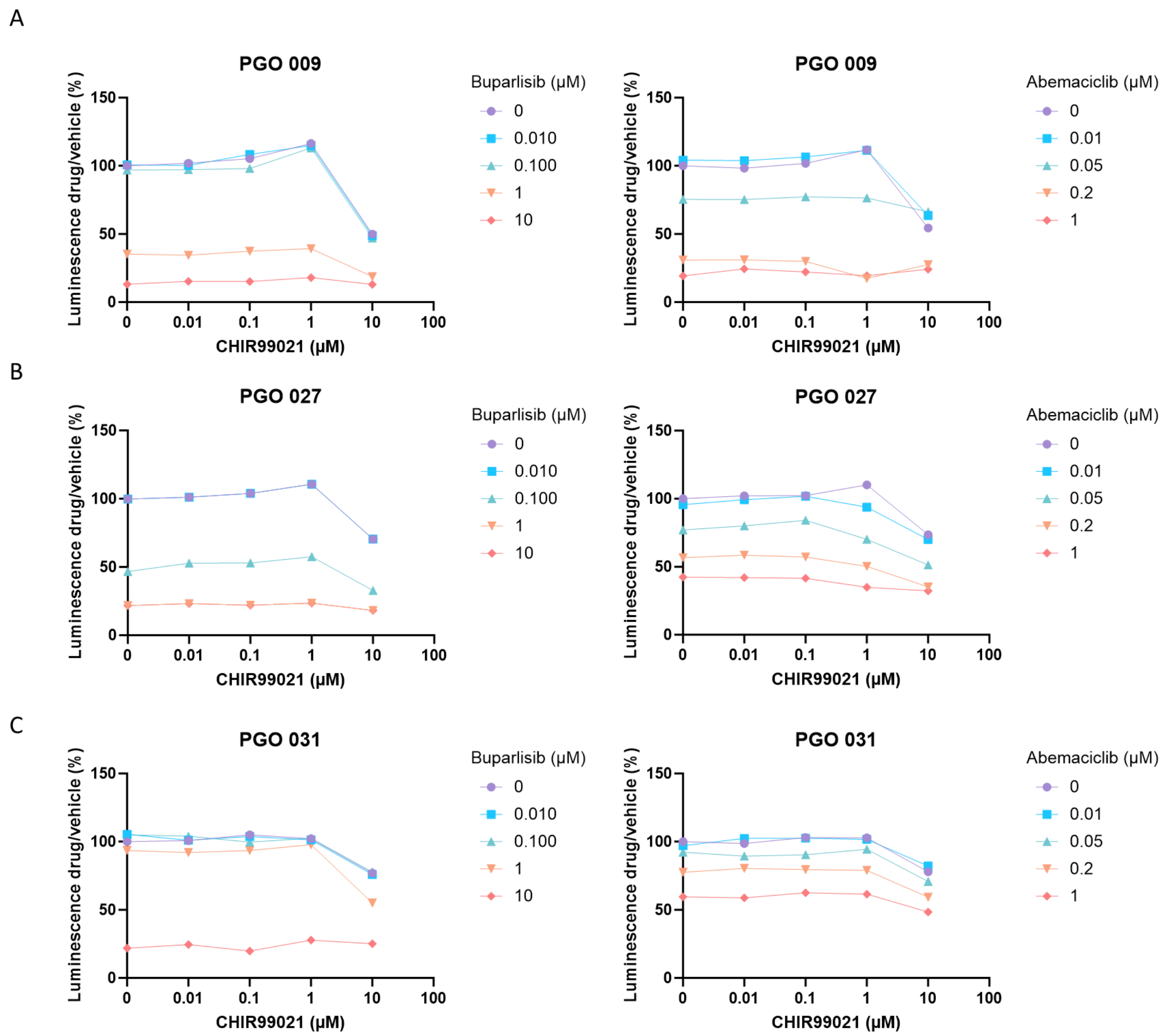

*Supplementary Figure 4: Cell viability of combinatorial treatment with CHIR99021 and buparlisib (A) or abemaciclib (B) in PGO 009, PGO 027 and PGO 031 (N=3)*

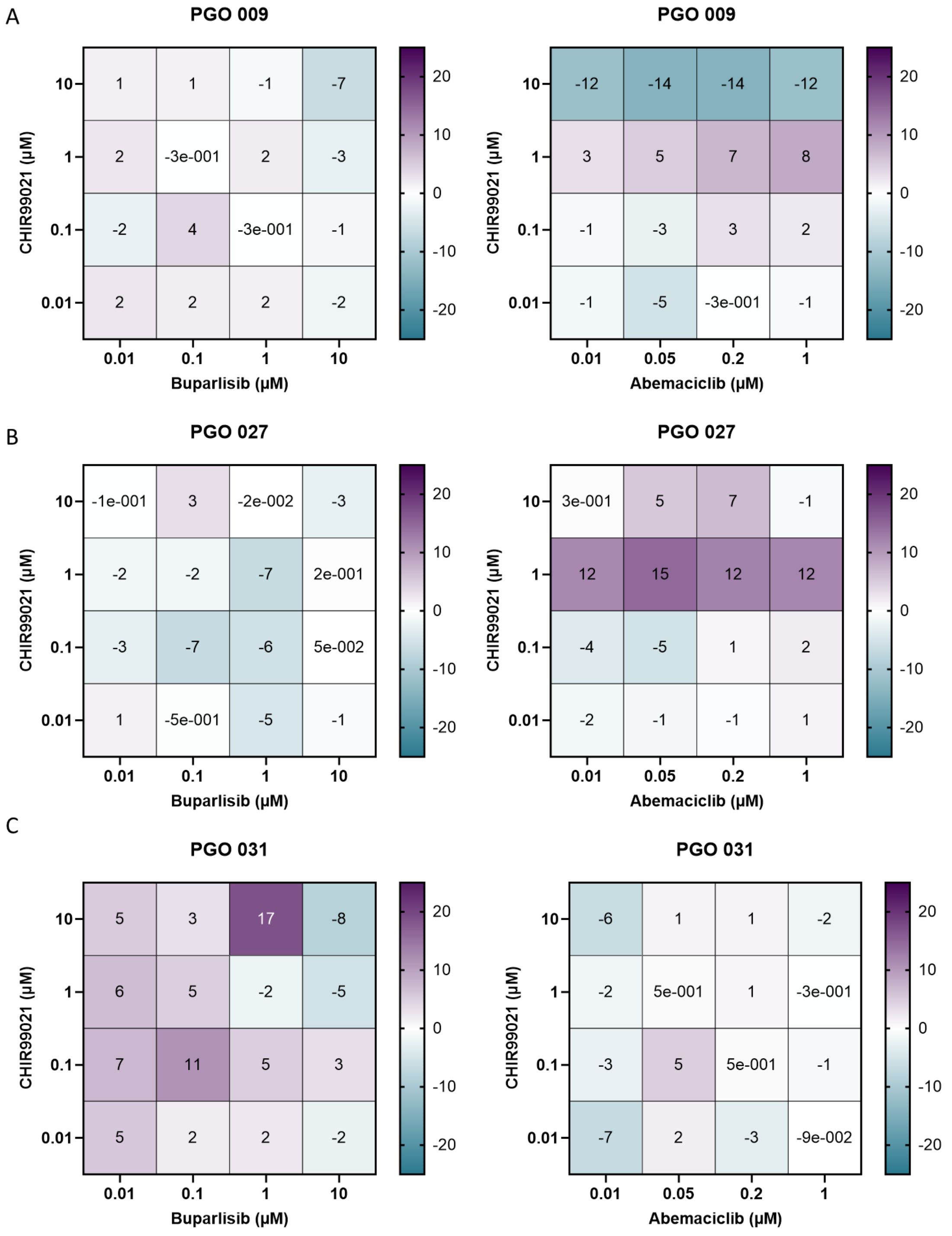

*Supplementary Figure 5: Bliss scores of combinatorial treatments with CHIR99021 and buparlisib (A) or abemaciclib (B) in PGO 009, PGO 027 and PGO31 (N = 3)*

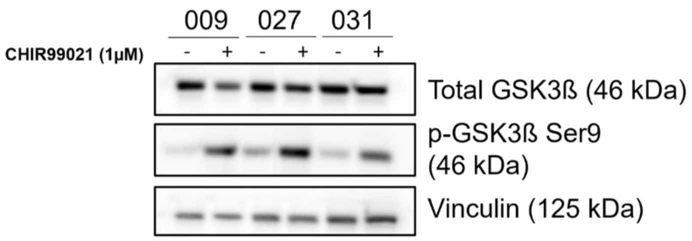

*Supplementary Figure 6: Western blot of on target efficacy of CHIR99021 after 24h treatment with 1µM CHIR99021 in PGO 009, PGO 027 and PGO 031 (N = 1).* Vinculin was used as a loading control.

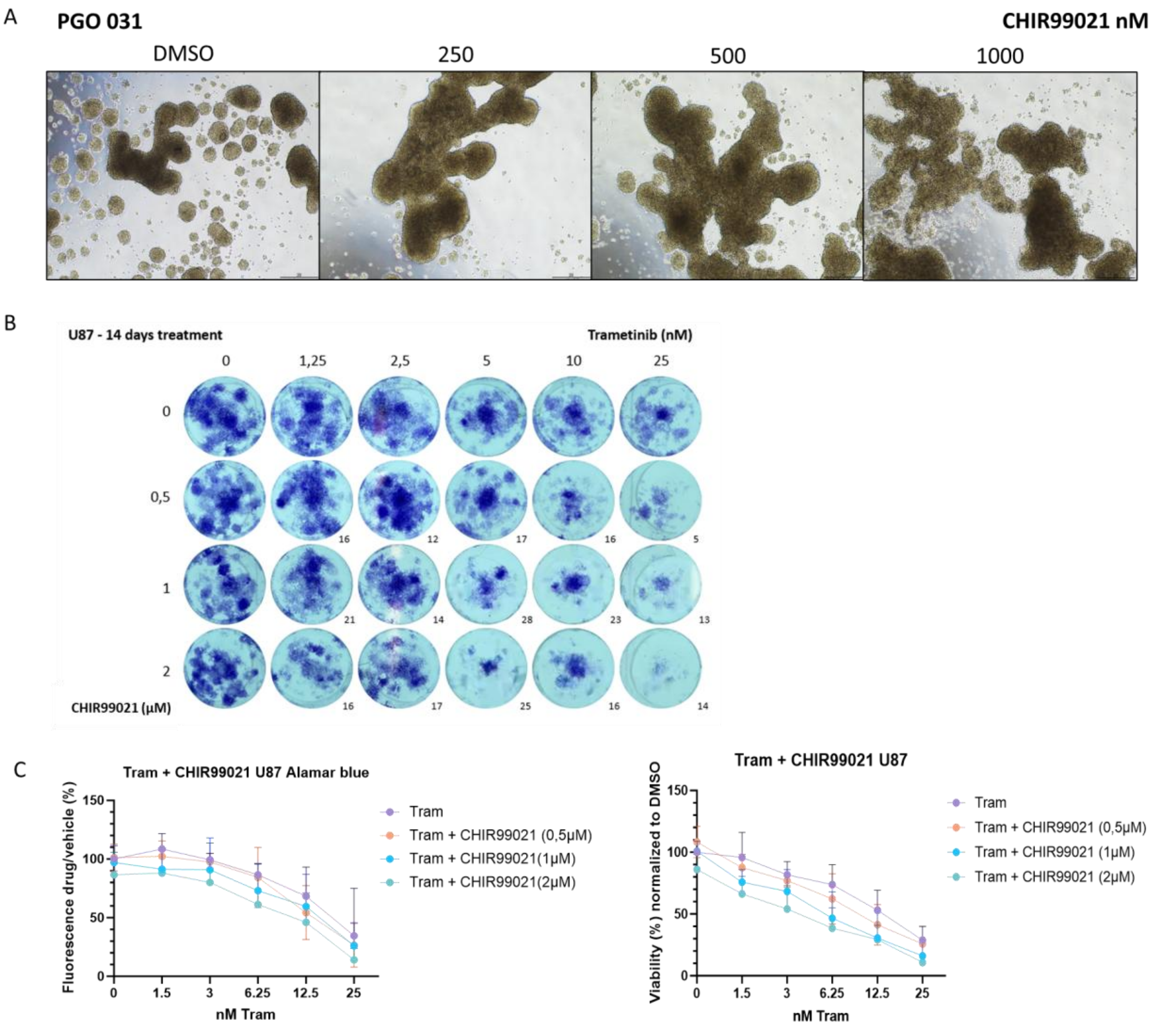

| Bliss scores | 1.5nM | 3nM | 6nM | 12.5nM | 25nM |
|---|---|---|---|---|---|
| 0,5µM | -12.5 | 0.6 | 3.4 | 14.7 | 8.4 |
| 1µM | 13.9 | 5.8 | 10.8 | 6.6 | 7.4 |
| 2µM | 5.9 | 6.1 | 13.6 | 13.1 | 15.7 |

| Bliss scores | 1.5nM | 3nM | 6nM | 12.5nM | 25nM |
|---|---|---|---|---|---|
| 0,5µM | -28.6 | 1.4 | 10.7 | 17.7 | 5.5 |
| 1µM | 24.3 | 22.3 | 16.6 | 16.6 | 7.1 |
| 2µM | 38.9 | 38.3 | 26.6 | 23.3 | 10.4 |

*Supplementary Figure 7: Long-term effects of CHIR99021 in combination with trametinib in PGO 031 and U87 GB cell lines*. (A) Brightfield microscopic images (20x) of PGO 031 after DMSO, 250, 500 and 1000 nM CHIR99021 (B) Crystal violet assay, where U87 has been exposed for 14 days. A single experiment is visualized from 4 biological replicates in triplo. Bliss scores have been given for combined treatment conditions. (C) Metabolic cell viability in U87 cells assessed using the Alamar Blue assay (n = 4). (D) Quantification of crystal violet staining intensity in U87 using Image Studio Lite (n = 4).

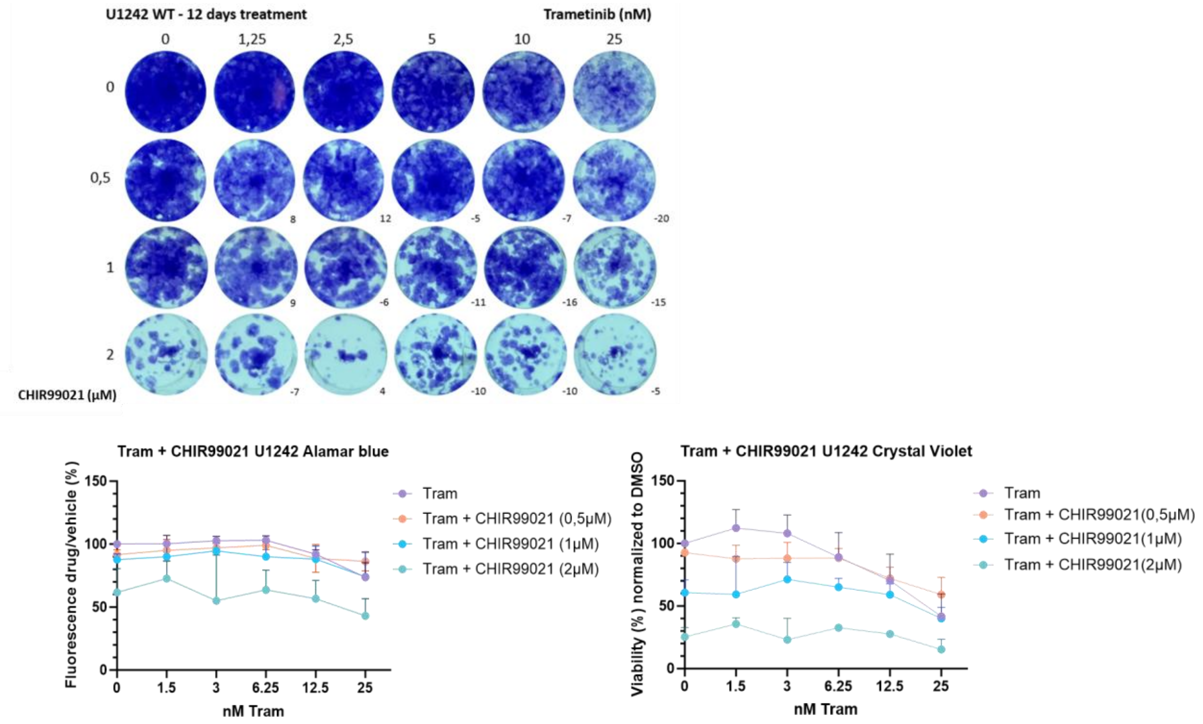

*Supplementary Figure 8: Long-term effects of CHIR99021 in combination with trametinib in U1242 GB cell lines.* (A) Crystal violet assay, where U1242 has been exposed for 14 days. A single experiment is visualized from 3 biological replicates in triplo. Bliss scores have been given for combined treatment conditions. (B) Metabolic cell viability in U1242 cells assessed using the Alamar Blue assay (n=3). (C) Quantification of crystal violet staining intensity in U1242 using Image Studio Lite (n=3).
